# Imputation-based fine-mapping suggests that most QTL in an outbred *chicken* Advanced Intercross Line are due to multiple, linked loci

**DOI:** 10.1101/061333

**Authors:** Monika Brandt, Muhammad Ahsan, Christa F. Honaker, Paul B. Siegel, Örjan Carlborg

## Abstract

The Virginia *chicken* lines have been divergently selected for juvenile body-weight for more than 50 generations. Today, the high-and low-weight lines show a 12-fold difference for the selected trait, 56-day body-weight. These lines provide unique opportunities to study the genetic architecture of long-term, single-trait selection. Previously, several Quantitative Trait Loci (QTL) contributing to weight differences between the lines were mapped in an F_2_-cross between them, and these were later replicated and fine-mapped in a nine-generation advanced intercross of them. Here, we explore the possibility to further increase the fine-mapping resolution of these QTL via a pedigree-based imputation strategy that aims to better capture the haplotype-diversity in the divergently selected, but outbred, founder lines. The founders of the intercross were high-density genotyped, and then pedigree-based imputation was used to assign genotypes throughout the pedigree. Imputation increased the marker-density 20-fold in the selected QTL, providing 6911 markers for the subsequent analysis. Both single-marker association and multi-marker backward-elimination analyses were used to detect associations to 56-day body-weight. The approach revealed several statistically and population-structure independent associations and increased the resolution of most QTL. Further, most QTL were also found to contain multiple independent associations, implying a complex underlying architecture due to the combined effects of multiple, linked loci on independent haplotypes that still segregate in the selected lines.

**Article summary:** After 50 generations of bi-directional selection, the Virginia chicken lines display a 12-fold difference in bodyweight at 56 days of age. Birds from the high and low selected lines were crossed to found an Advanced Intercross Line, which has been maintained for 9 generations. Using high-density genotypes of the founders, we imputed genotypes in intercross birds that were only genotyped for a sparse set of markers. Using single and multi-marker association analyses, we replicated nine known body-weight QTL. Multiple statistically independent associations were revealed in eight of the QTL, suggesting that most are caused by multiple linked loci.

## Introduction

Long-term selective breeding of animals and plants for extreme phenotypes has resulted in genetically distinct lines that are a valuable resource for dissecting the genetic architecture of complex traits (Hill 2005). Most traits of interest in animal breeding (e.g. production of eggs or meat, resistance to disease) are influenced by a combination of genetic and environmental factors. Due to their multi-factorial nature and despite the ability to obtain data on both genome-wide genetic markers and phenotypes from large numbers of individuals, it is challenging to disentangle their genetic architecture when working with commercial populations. An alternate strategy is to make use of experimental populations resulting from long-term selection experiments, where the focus has been to develop divergent lines from a common base-population using more coherent selection criteria. Such populations will display larger phenotypic differences than populations subjected to composite, commercial breeding programs and hence facilitate in-depth studies of the genetic basis underlying the selection-response and general genetic architecture of these traits (Andersson and Georges 2004). Given that many of the agriculturally important traits relate to metabolism, feeding-behavior and growth, they also provide a good model for translational studies to decipher the genetic architecture of traits of interest in human medicine, including obesity, eating disorders, and diabetes (Andersson 2001).

The Virginia lines are experimental populations established in 1957 to study the genetic effects of long-term (more than 50 generations), divergent, single-trait selection for 56-day high (HWS) or low (LWS) body-weight in *chickens* (Dunnington and Siegel 1996; Márquez *et al.* 2010; Dunnington *et al.* 2013). The lines originated from the same base population, composed by crossing seven partially inbred White Plymouth Rock *chicken* lines, and today display a 12-fold difference in body-weight at 56 days of age (Márquez *et al.* 2010; Dunnington *et al.* 2013). In addition to the direct effects of selection on body-weight, the selected lines also display correlated selection responses for a range of metabolic and behavioral traits including disrupted appetite, obesity, and antibody response (Dunnington *et al.* 2013).

The Virginia HWS and LWS lines have been used extensively for studying the genetic architecture of body-weight and other metabolic traits. These studies have uncovered a number of loci with minor direct effects on body-weight, metabolic traits and body-stature traits by Quantitative Trait Loci (QTL) mapping in an F_2_ intercross (Jacobsson *et al.* 2005; Park *et al.* 2006; Wahlberg *et al.* 2009). Also, a network of epistatic loci has been found to make a significant contribution to long-term selection response through the release of selection induced additive variation (Carlborg *et al.* 2006; Le Rouzic *et al.* 2007; Le Rouzic and Carlborg 2008). Explorations of the genome-wide footprint of selection by selective-sweep mapping suggests that perhaps more than one hundred loci throughout the genome have contributed to selection response (Johansson *et al.* 2010; Pettersson *et al.* 2013), and many of these contribute to 56-day body weight (Sheng *et al.* 2015).

To replicate and fine-map the body-weight QTL inferred in the F_2_ intercross, we developed, genotyped and phenotyped, for body-weight at 56 days of age (BW56), a nine-generation Advanced Intercross Line (AIL). This large AIL originated from the same founders as the F2 intercross, but was selectively genotyped at a higher resolution (~1 marker/cM) in nine QTL (Besnier *et al.* 2011). In this population, most of the original minor (Besnier *et al.* 2011) and epistatic (Pettersson *et al.* 2011) QTL were replicated and fine-mapped. These earlier studies analyzed the data using a haplotype-based linkage-mapping approach in a variance-component based model framework to infer single-locus effects (Besnier *et al.* 2011) or a fixed-effect model framework assuming fixed alternative alleles in the two founder-lines for detecting epistasis (Pettersson *et al.* 2011). The variance-component model was used in the replication study in order to avoid the assumption of allelic fixation in the founder-lines. By implementing it in a Flexible Intercross Analysis (FIA) modeling framework (Rönnegård *et al.* 2008), it was expected to improve power when the parental lines carry alleles with correlated effects (e.g. multiple alleles with similar effects).

Although the initial studies mapped QTL under the assumption of fixation, or an effect correlation, of divergent alleles in the crossed lines, the results at the same time implied that multiple alleles might be segregating in several of the mapped regions. To this end, the first QTL replication study in the AIL population (Besnier *et al.* 2011) found a large within founder-line heterogeneity in the allelic effects. Later the selective-sweep studies, that utilized data from multiple generations of divergently selected and relaxed lines, identified on-going selection and multiple sweeps in many QTL (Johansson *et al.* 2010; Pettersson *et al.* 2013), as well as extensive allelic purging (Pettersson et al. 2013). This allelic heterogeneity challenges attempts to dissect the architecture of the selected trait via e.g. QTL introgression (Ek *et al.* 2012). Alternative approaches are therefore needed to uncover multi-locus, multi-allelic genetic architectures in QTL and their contributions to the long-term response to directional selection.

In this study, to explore an imputation-based association-mapping strategy for further dissection of previously mapped and replicated QTL (Besnier *et al.* 2011; Pettersson *et al.* 2011), we made use of available high-density (60K SNP-Chip) genotypes for founders (Johansson *et al.* 2010; Pettersson *et al.* 2013) and intermediate-density SNP-genotypes in several QTL in the entire 9-generation AIL-pedigree. By increasing the marker-density in the QTL throughout the AIL by imputation, we aimed to better capture the segregating haplotypes within and between the divergently selected founder populations than with the previously used markers. This aim can be achieved as the original markers genotyped in the AIL were selected to identify high-and low-line derived alleles, and not alleles that segregate within or across the founder lines. By testing for association between imputed markers and body-weight, the fine-mapping analyses were less constrained by the original selection of markers and facilitated a more thorough exploration of the genetic architectures of the nine evaluated QTL. Our results show that the imputation-based approach not only allows replication of most QTL, but also that it is possible to utilize historical recombination in the pedigree to improve the resolution in the fine-mapping analyses. We found that several of the original QTL are likely due to the combined effects of multiple linked loci, several of which are segregating for alleles originating from different haplotypes in the founder population of the selected lines.

## Materials and methods

### Animals

The Virginia *chicken* lines are part of an ongoing selection experiment to study the genetics of long-term, single-trait selection (Márquez *et al.* 2010; Dunnington *et al.* 2013). It was initiated in 1957 from a base population, generated by intercrossing seven partially inbred lines of White Plymouth Rock *chickens*. From the offspring of the partially inbred lines, resulting from the intercrossing, the birds with the highest and lowest 56-day body-weights (with some restrictions), respectively, were selected to produce the high-and low-weight selected lines (HWS and LWS) (Márquez *et al.* 2010; Dunnington *et al.* 2013). Since then, the lines have undergone divergent selection for increased and decreased body weights with one new generation hatched in March of every year.

An Advanced Intercross Line (AIL) was founded by reciprocal crosses of 29 HWS and 30 LWS founder birds from generation 40 (Besnier *et al.* 2011). The mean, sex-averaged 56-day body weights for HWS and LWS at this generation were 1522 g and 181 g, respectively. Repeated intercrossing of birds was used to develop a nine-generation AIL consisting of generations F_0_-F_8_. In each generation, approximately 90 birds were bred by paired mating, genotyped, and weighed at 56 days of age (BW56). The exceptions were generations F_3_ and F_8_ that contained 405 and 437 birds, respectively (Besnier *et al.* 2011). In total, the AIL population consisted of 1529 F_0_ to F_8_ individuals with complete records on pedigree and genotypes (see below), and 1348 F_2_-F_8_ individuals with juvenile body-weight (BW56) records.

### Genotyping

The complete AIL pedigree (1529 birds) had earlier been genotyped in nine selected QTL for 304 SNP-markers that passed quality-control as described in (Besnier *et al.* 2011). Further, 40 of the founders for the pedigree (20 HWS and 20 LWS) had also earlier been genotyped using a whole genome 60K SNP chip (Johansson *et al.* 2010; Pettersson *et al.* 2013). The 6607 markers from the SNP-chip that were informative and passed quality control in that study are located in the nine QTL-regions targeted in this study. When merging the information from the 60K SNP chip and the information from the 304 markers genotyped earlier, 55 markers in 40 founders were genotyped using both methods. Out of these 55 markers, 28 markers with genotype inconsistencies between the genotyping technologies were removed during quality control. In total, our analyses were based on 6888 markers, where 40 of the 59 AIL founders had genotypes for all markers, and the remaining individuals in the pedigree had genotypes for 281 markers. Table 1 shows how these markers are distributed across the nine QTL regions.

**Table 1.**
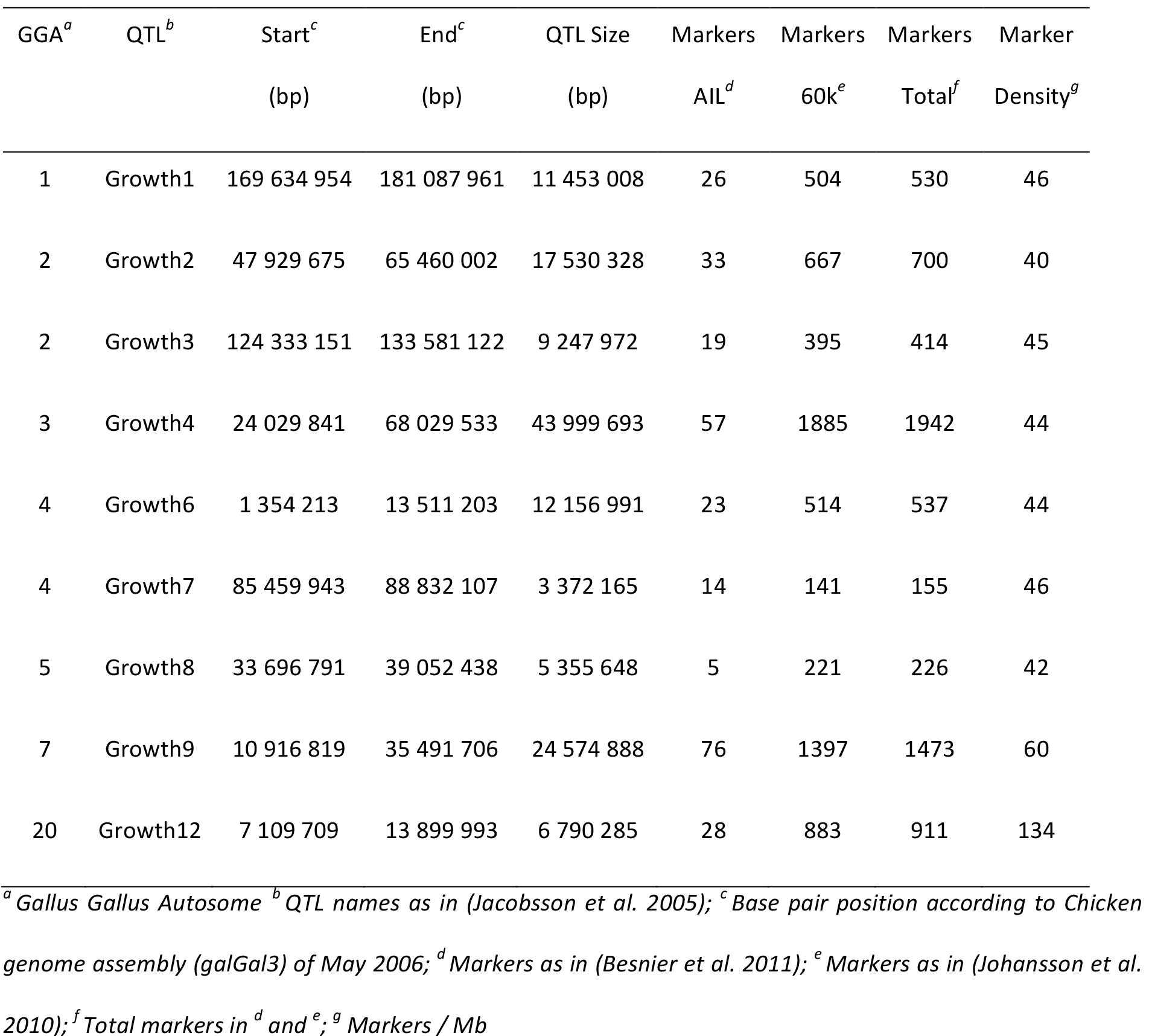
Genotyped and imputed markers across the nine analyzed QTL.

| GGA<sup>a</sup> | QTL<sup>b</sup> | Start<sup>c</sup> | End<sup>c</sup> | QTL Size | Markers | Markers | Markers | Marker |
| --- | --- | --- | --- | --- | --- | --- | --- | --- |
|  |  | (bp) | (bp) | (bp) | AIL<sup>d</sup> | 60k<sup>e</sup> | Total<sup>f</sup> | Density<sup>g</sup> |
| 1 | Growth1 | 169 634 954 | 181 087 961 | 11 453 008 | 26 | 504 | 530 | 46 |
| 2 | Growth2 | 47 929 675 | 65 460 002 | 17 530 328 | 33 | 667 | 700 | 40 |
| 2 | Growth3 | 124 333 151 | 133 581 122 | 9 247 972 | 19 | 395 | 414 | 45 |
| 3 | Growth4 | 24 029 841 | 68 029 533 | 43 999 693 | 57 | 1885 | 1942 | 44 |
| 4 | Growth6 | 1 354 213 | 13 511 203 | 12 156 991 | 23 | 514 | 537 | 44 |
| 4 | Growth7 | 85 459 943 | 88 832 107 | 3 372 165 | 14 | 141 | 155 | 46 |
| 5 | Growth8 | 33 696 791 | 39 052 438 | 5 355 648 | 5 | 221 | 226 | 42 |
| 7 | Growth9 | 10 916 819 | 35 491 706 | 24 574 888 | 76 | 1397 | 1473 | 60 |
| 20 | Growth12 | 7 109 709 | 13 899 993 | 6 790 285 | 28 | 883 | 911 | 134 |

<sup>a</sup> *Gallus Gallus* Autosome <sup>b</sup> QTL names as in (Jacobsson *et al.* 2005); <sup>c</sup> Base pair position according to Chicken genome assembly (*galGal3*) of May 2006; <sup>d</sup> Markers as in (Besnier *et al.* 2011); <sup>e</sup> Markers as in (Johansson *et al.* 2010); <sup>f</sup> Total markers in <sup>d</sup> and <sup>e</sup>; <sup>g</sup> Markers / Mb

### Phasing and imputation of markers

All genotyped markers in the QTL (Table 1) were first ordered according to their physical location in the *Chicken* genome assembly of May 2006 (*galGal3*). In the ordered marker set, the SNP-chip markers were evenly distributed in the intervals between the sparser set of markers genotyped across the entire AIL.

Using the software ChromoPhase (Daetwyler *et al.* 2011), we phased and imputed genotypes for the complete set of 6888 markers across the entire AIL pedigree. ChromoPhase first phases large segments of chromosomes, in our case the QTL regions. It then imputes the missing genotypes in the AIL individuals genotyped with the sparse set of markers from the genotype information available in high-density genotyped founders utilizing the pedigree information. It thus predicts both phased haplotypes across the nine studied QTL and genotypes at markers that were only genotyped in a subset of the founder individuals in the pedigree.

### Single-marker association analyses

The *qtscore* function in the GenABEL package (Aulchenko *et al.* 2007) was used to test for association between body-weight at 56 days of age and, genotyped or imputed, individual genetic markers within the targeted QTL. The allelic effect of each marker, 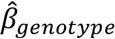, was estimated using a regression model (Model 1), where the genotype at each marker was coded in ***Z*** as 0 if homozygous for the major allele, 1 if heterozygous, and 2 if homozygous for the minor allele. Sex and generation was added as categorical covariates, with 2 different levels for sex and 7 different levels for generation, defined for each individual in ***X***. The phenotype, body-weight at 56 days of age, is given in the numerical variable **y**.

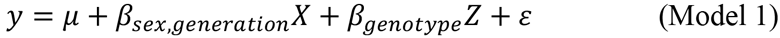

*ɛ* was assumed to be iid and normally distributed around 0 with variance *σ*^2^. *μ* is the intercept, that in this model represented the mean body-weight at 56 days of age for female individuals from the F_2_ generation.

### Multi-locus association analysis accounting for population-structure

The single-marker association analyses based on Model 1 neither account for possible genetic dependencies (linkage or LD) between markers within the QTL regions, nor account for the potential effects of population-structure in the AIL (Peirce *et al.* 2008; Cheng *et al.* 2010). Earlier studies have shown that the genetic architecture of body-weight is highly polygenic in this population (Jacobsson *et al.* 2005; Wahlberg *et al.* 2009; Johansson *et al.* 2010; Besnier *et al.* 2011; Pettersson *et al.* 2011; 2013; Sheng *et al.* 2015) and we therefore implemented a statistical approach based on this. To correct for population-structure in the deep intercross line and identify an experiment-wide set of independent association-signals, we use a forward-selection/backward-elimination procedure in a bootstrap-based framework developed for this purpose (Valdar *et al.* 2009; Sheng *et al.* 2015). As all markers with genotypes could not be included in a backward-elimination analysis due to the limited sample size, we first used a forward-selection strategy to identify a smaller set of statistically suggestive independent signals within each QTL region to be thoroughly evaluated in the backward-elimination analysis (Valdar *et al.* 2009; Sheng *et al.* 2015). The forward-selection analysis was performed by scanning across all markers within each QTL using Model 1. If any of the markers were nominally significant (p < 0.05) in the scan, the marker with the strongest association was added as a covariate in the model. This procedure was repeated until no more significant markers were detected. The models were fitted using the *qtscore* function in the *GenABEL* package (Aulchenko *et al.* 2007) as described previously. The markers from this analysis with an allele-frequency > 0.10 in the population were subjected to the full backward-elimination analysis described below.

In short, we used a bootstrap based backward-elimination model-selection framework (Sheng *et al.* 2015) across the markers selected by forward-selection in the QTL. An adaptive model selection criterion controlling the False Discovery Rate (Abramovich *et al.* 2006; Gavrilov *et al.* 2009) was used during backward-elimination in a standard linear model framework, starting with a full model including the fixed effects of sex and generation, and the additive effects of all markers (Model 2).:

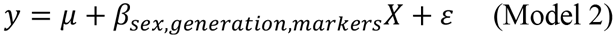

where phenotype, sex, and generation are coded as described for model 1 and where *ɛ* again is assumed to be iid and normally distributed around 0 with variance *σ*^2^. The intercept, *μ*, represents the mean body-weight at 56 days of age for female individuals from the F_2_ generation. In model 2, genotypes are coded based on the line-origin of the alleles at each locus. If an individual was homozygous for the major allele in the AIL-founders from the high-weight selection line, its genotype was coded as 1 at that locus. If an individual was heterozygous, its genotype was coded as 0. If an individual was homozygous for the allele corresponding to the major allele in AIL-founders from the low-weight selection line, its genotype was coded as −1. By coding genotypes in a −1, 0, and 1 manner, the estimates, 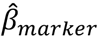, from fitting model 2 will be negative if a weight-decreasing allele is derived from the high-weight line or if a weight-increasing allele is derived from the low-weight line.

Convergence was based on a 20% FDR level. The analysis was performed using bootstrapping with 1000 resamples. Markers with an RMIP (Resample Model Inclusion Probability) > 0.46, as suggested for an AIL generation F_18_ (Valdar *et al.* 2009), was included in the final model. The FDR in the final model was confirmed using the original FDR procedure described in (Benjamini and Hochberg 1995) as implemented in the p.adjust function in the R stats-package (R Development Core Team 2015). The additive genetic effect for each locus was estimated using the multi-locus genetic model described above (Model 2).

### Data availability

Genotype and pedigree data will be included as supplementary information in the published version of the manuscript.

## Results and Discussion

We compared the results of the imputation-based association analyses with the previously reported results from the linkage-based analysis of the same nine QTL in Besnier *et al.* (2011). Figure 1 shows the statistical support for association and linkage to BW56 across the QTL. As the imputation-based analysis does not model pedigree-relationships, comparisons were made to the results from model A in (Besnier *et al.* 2011). Overall, the results between the two analyses overlap well. For several of the QTLs, we were able to fine-map the associated regions using imputed genotypes for SNPs within these regions. The overall results from the single-marker association analyses also agree with those from the bootstrap-based forward-selection/backward-elimination approach used to identify genome-wide independent association signals.

### Four statistically independent associated markers in the GGA7 QTL *Growth9*

The QTL *Growth9* on GGA7 (Gallus Gallus Autosome 7) (Figure 1; 10.9-35.5 Mb) was the only QTL that reached genome-wide significance in the first F_2_ intercross between the HWS and LWS lines (Jacobsson *et al.* 2005). It was later identified as a central locus in an epistatic network explaining a large part of the difference in weight between HWS and LWS lines (Carlborg *et al.* 2006). In the earlier fine-mapping analysis, the linkage signal covers most of the QTL region (from 15-35 Mb), but further analyses showed that two independent loci were segregating in the region (Besnier *et al.* 2011). The signal in the imputation-based association analysis performed here is more focused, with a highly significant signal in a 2.8 Mb region between 23.7-26.4 Mb that overlaps with the strongest signal in the linkage-scan. Previously, Ahsan *et al.* (2013) explored potential candidate mutations in the QTL and found two regulatory SNPs near the peak at 21 Mb (21.6 and 22.7 Mb) and a synonymous-coding SNP in a CpG island in an exon of the Insulin-like growth factor binding protein 2 (*IGFBP2*) gene in the middle of the major association peak at 24.8 Mb. In addition to the strong association around 24 Mb, the association analysis also highlights two additional regions (centered around 18 Mb and 29 Mb) that are also shown to be experiment-wide independent signals in the backward-elimination analysis. The second QTL detected in the linkage analysis (*Growth9.2*) was only nominally significant in the single-marker association analysis, but as a marker in this region was included in the final model from the bootstrap-based backward-elimination analysis this suggests that the effect of this locus might be dependent on the genotype at other loci. In Table 2, we provide the additive effects and the significance of the experiment-wide independent loci from the backward-elimination analysis.

**Figure 1.**
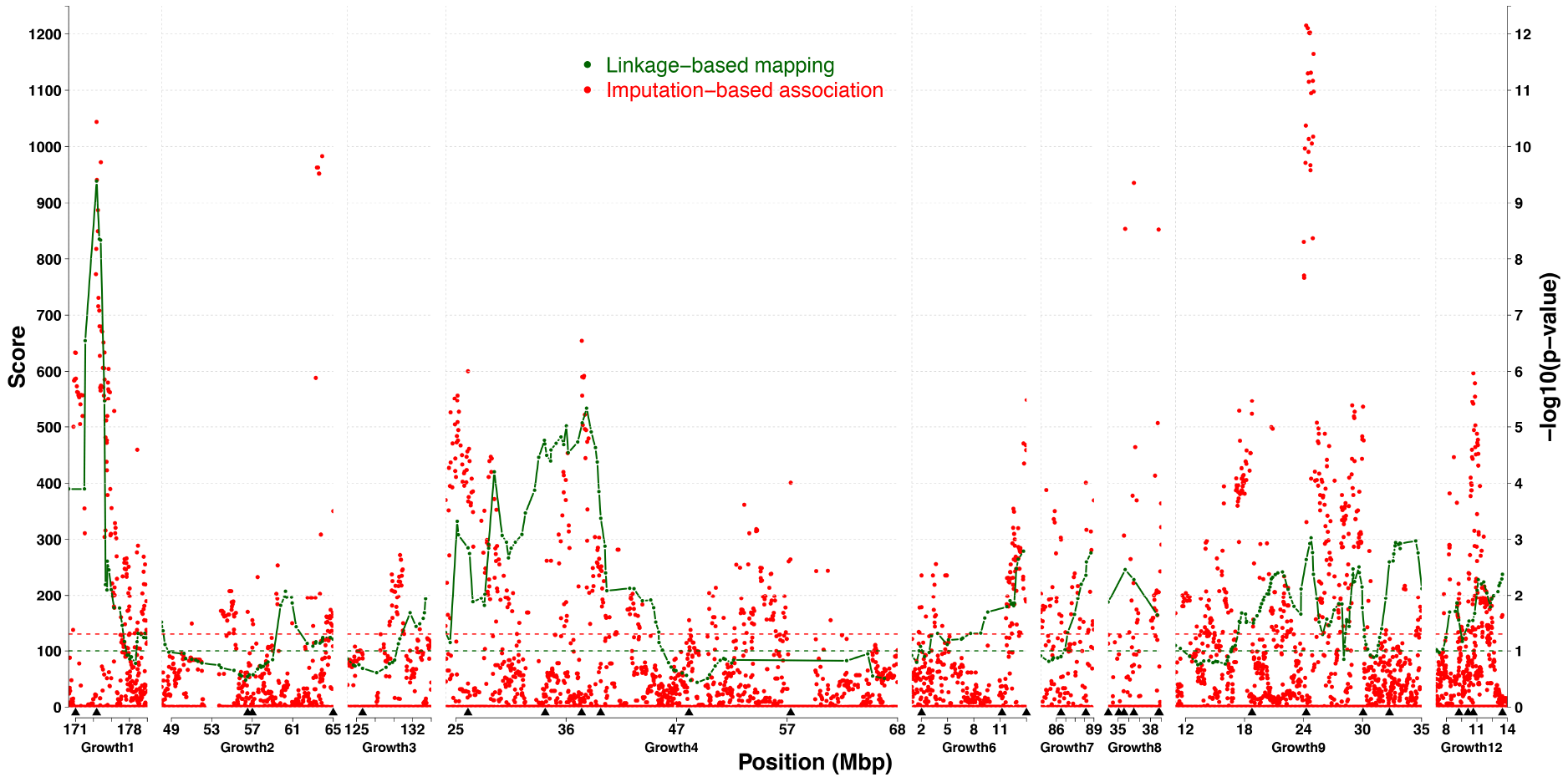
Comparison between linkage and association-based fine-mapping analyses of nine QTL in an Advanced Intercross Line chicken population. Green lines show the statistical support curve (score statistics from model A) for the linkage-based mapping study of Besnier et al. (2011) and the red dots associations to each analyzed marker in the new imputation-based association analysis (this study). The green and red horizontal dotted lines indicates the significance thresholds for the linkage-analysis threshold and the nominal significance threshold in the imputation-based association analysis, respectively. Arrowheads under the x-axis indicate the position of markers identified as experiment-wide significant in the bootstrap-based backward-elimination procedure.

**Table 2.**
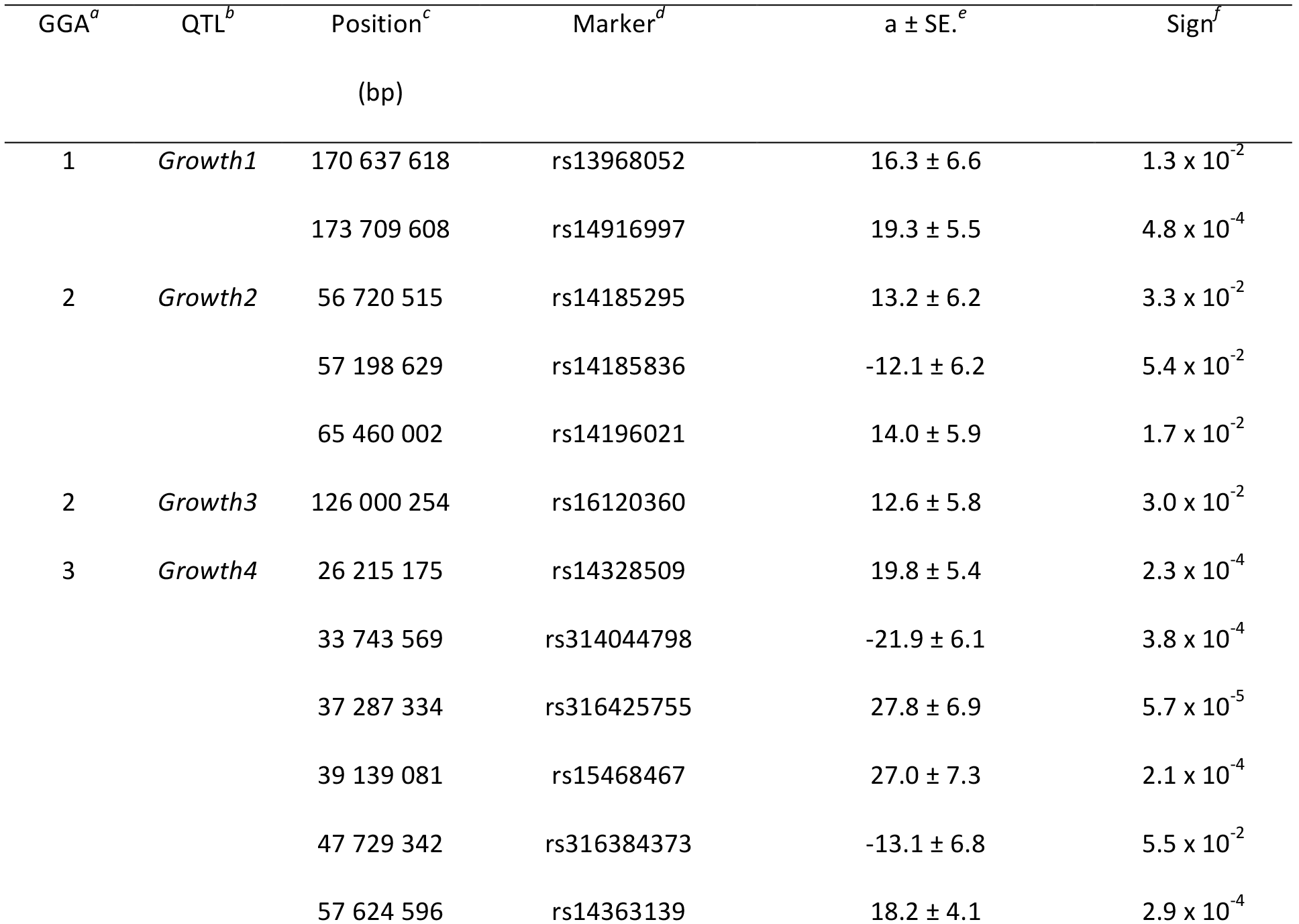
Estimated additive effect and standard error for experiment-wide independent association signals, between body-weight at 56 days of age and genotype, identified in a bootstrap based approach implemented in a backward-elimination model-selection framework across the markers in the genotyped QTL. For a marker with a positive estimated additive effect, the effect on weight is caused by the allele with its origin in the line associated with the sign of the effect, i.e. an allele with its origin in high-line is associated with an increase in body-weight and an allele with its origin in low-line is associated with a decrease in body-weight. In cases where a weight-increasing allele has its origin in the low-line or a weight-decreasing allele has its origin in the high-line the sign of the estimated additive effect will be negative.

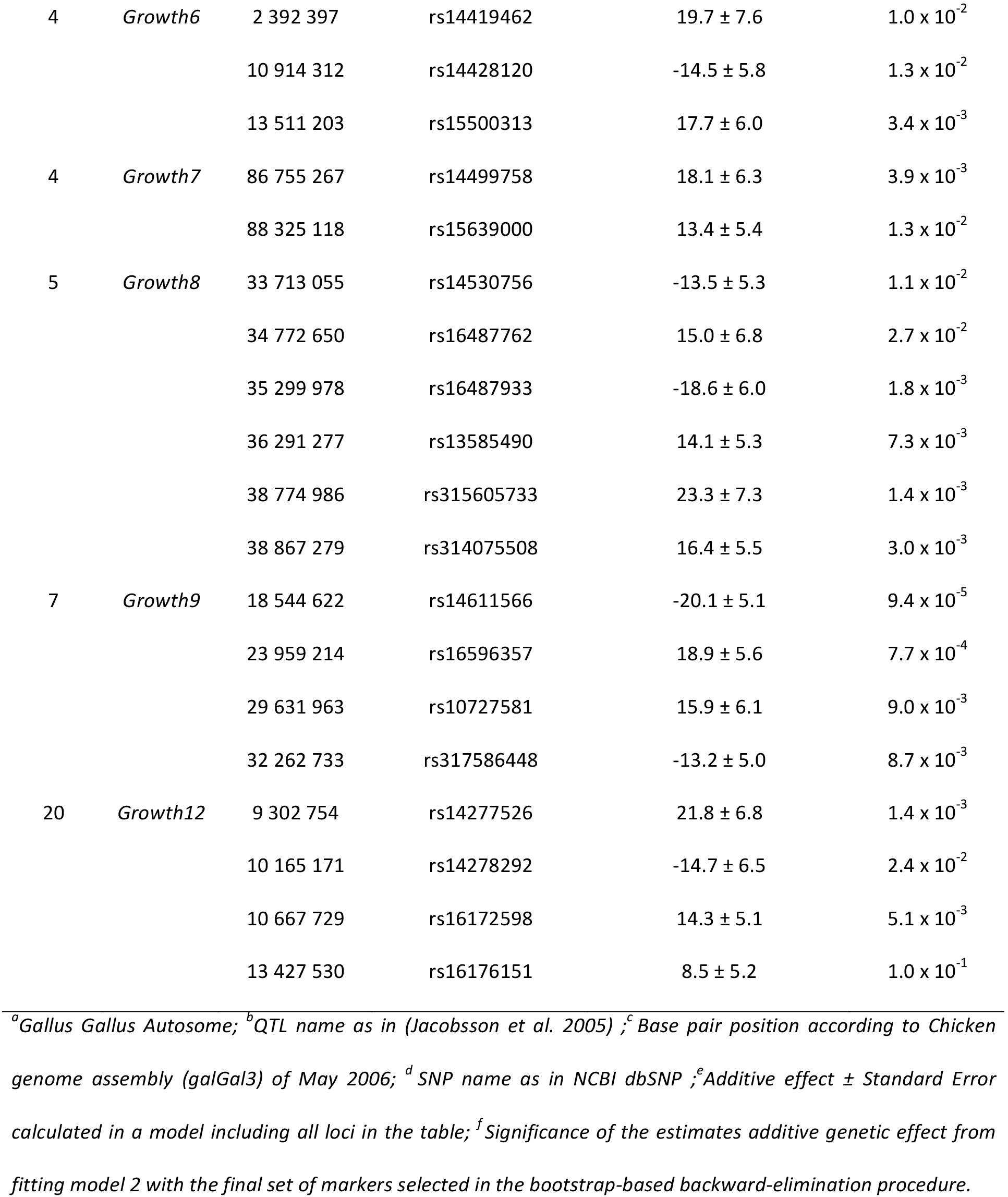

### Two statistically independent associated markers in the GGA1 QTL *Growthl*

The strongest association in the study by Besnier *et al.* (2011) was found on GGA1 in the QTL *Growth1* (Figure 1; 169.6-181.1 Mb). Here, the second strongest association was detected in that QTL. The imputation-based association analysis, however, highlights two significant associations, separated by a region of very low association. The strongest of these association-peaks was located near the peak detected using the earlier linkage-based analysis. Several of the significant associated markers were located in this region (173.6 to 175.3 Mb). A candidate gene for growth, Asparagine-linked glycosylation 11 homolog gene (*ALG11*), is located at 174.6 Mb and has a strong mutation in its regulatory region (Ahsan et al. 2013). The second association was found to a group of significant markers in a narrower region upstream from the main linkage-peak (170.3 and 171.7 Mb). The association analysis thus suggests that the original 10.6 Mb QTL region is due to the effects of two separate loci located in these confined 1.5 Mb and 1.8 Mb regions. This is further supported by the forward-selection/backward-elimination procedure, which identifies two experiment-wide independent signals to markers within *Growth1*, one at 170.6 Mb and one at 173.7 Mb.

### Four statistically independent associated markers in the GGA2 QTLs *Growth2 and Growth3*

GGA2 contain two QTL. *Growth2* (Figure 1; 47.9-65.5 Mb) has one highly associated peak at 64.3 Mb. *Growth3* (Figure 1; 124.3-133.6 Mb) was the least significant QTL in the association analysis, with a small peak at 129 Mb. The forward-selection/backward-elimination analyses identify 3 significant associations in *Growth2* at 56.7 Mb, 57.2 Mb, and 65.5 Mb respectively, with the strongest significance at 65.5 Mb. This suggests that there are two distinct associated loci in *Growth2*. In the earlier linkage-based analysis the strongest signal in *Growth2* was found at 60.6 Mb. In *Growth3* the strongest signal detected here is shifted almost 4 Mb upstream from the top signal found in the earlier linkage-based analysis.

### Six statistically independent associated markers in the GGA5 QTL *Growth8*

One of the strongest association signals was found on GGA5 in the QTL *Growth8* (Figure 1; 33.7-39.1 Mb). The most associated markers were located in the central part of this 5.3 Mb QTL and overlaps with the earlier linkage-signal. The association signal was, however, stronger than the linkage signal suggesting that the imputed markers tag the QTL better than the haplotypes inferred from the sparser set of genotyped markers. The results from the forward-selection/backward-elimination analyses (Table 2) suggest that several markers tag the variation present in this region and the strongest associated SNP was located at 38.7 Mb.

### Six statistically independent associated markers in the GGA3 QTL *Growth4*

In the QTL *Growth4* on GGA3 (Figure 1; 24.0-68.1 Mb), both the association and linkage analyses find the strongest signals between 24-41 Mb. Although the statistical support curve in the linkage-analysis contains multiple peaks, that analysis was unable to fine-map the region into multiple, independent signals. The association-analysis, however, revealed at least two distinct association-peaks with several associated markers in each region. These regions are located approximately 10 Mb apart and are separated by a region with very low association. The regions are mapped with high resolution and are located approximately between 24-27 Mb and 34-37 Mb, respectively. A candidate mutation in *Growth4* was found slightly outside the second association region at 33.6 Mb inside the regulatory region of Cystein rich transmembrane BMP regulator 1 (*CRIM1*) (Ahsan *et al.* 2013). In addition to the two peaks described above, several nominally significant markers were located around 55-57 Mb, suggesting that also this region is associated with BW56. This is supported by the forward-selection/backward-elimination analyses where the highly associated marker at 57.6 Mb is defined as experiment-wide significant. In the previously performed linkage-based analysis, this region displayed very low significance.

### Four statistically independent associated markers in the GGA20 QTL *Growth12*

The earlier linkage-analysis replicated the QTL *Growth12* on GGA20 (Figure 1: 7.1-13.9 Mb), with the strongest associated marker at 10.7 Mb, and the signal covered most of the region (8-13.9 Mb). In the association-analysis, several markers reached the nominal threshold and these are part of a focused association peak covering the region around 9 Mb and the region around 11 Mb. Both regions were identified as independent peaks in the backward-elimination analysis, suggesting that this QTL contains multiple associated loci.

### Five statistically independent associated markers in the GGA4 QTLs *Growth6 and Growth7*

In both *Growth6* (Figure 1; 1.3-13.6 Mb) and *Growth7* (Figure 1.; 85.4-88.9 Mb) on GGA4, several markers were nominally significant in the association analysis. These markers were located very close to the main peaks in the earlier linkage-based analysis, suggesting that the two analyses identify the same underlying loci. The association analysis highlighted a region in *Growth7* with strong association around 86 Mb that was not found with the linkage-based approach. As for *Growth12*, this suggests that the *Growth7* also contains multiple associated loci.

### General comments

Here we report the results from using an imputation-based association-mapping strategy to fine-map QTL in a nine-generation, outbred Advanced Intercross Line (AIL). By combining high-density genotyping of the AIL founders with imputation throughout the rest of the pedigree utilizing a sparser genotyped marker-backbone, we increased the marker-density ~20-fold in the studied regions. This subsequent association analysis had a comparable power for replication of QTL to the earlier used linkage-based strategy. In addition to this, the new analyses also detected multiple association-peaks in several of the QTL and narrowed the associated regions considerably compared to the regions detected previously (Besnier *et al.* 2011). Together, they suggest that this imputation-based association-mapping approach is a promising strategy for improving the resolution in fine-mapping studies in outbred pedigrees, where high-density marker genotypes are not available for all studied individuals.

In both *Growth1* and *Growth4* two strong, distinct association signals were identified. Also in the QTL *Growth8* and *Growth9* the new analysis identified strong association-peaks covering many markers. In these regions, the strongest linkage-signals identified in the previous fine-mapping analysis (Besnier *et al.* 2011) overlap with the strongest signals in the current analyses. However, the association analysis also separates the signals into multiple peaks and highlights narrower regions. Hence, it provides more useful input for further analyses to identify candidate genes underlying the QTL. In most cases the associated regions are restricted to distinct 2-3 Mb regions, which as indicated by the findings from Ahsan *et al.* (2013), is useful for restricting the bioinformatics analyses to only the most promising candidate genes for further functional studies. The additive effect of the marker identified as experiment-wide significant at 18.5 Mb in *Growth9* was assessed as transgressive. This, together with the extended linkage signal and the problem to replicate the QTL through introgression (Ek *et al.* 2012) suggests that the genetic architecture of the original *Growth9* QTL is more complex than previously noted, potentially due to the effects of more than two linked loci.

In *Growth6, Growth7*, and *Growth12*, the association signals were not as significant as in the other QTL. Despite this, the multi-locus analyses suggest that the linkage signals in the earlier analyses were due to distinct loci with independent effects, mapped here into narrower association peaks.

Overall, the location of the association-signals in this study overlapped well with the top signals in the earlier linkage analyses. However, in two of the QTL (*Growth2* and *Growth3*), the association peaks are shifted when comparing results from the two studies. Further work is needed to explore whether this reflects separate loci with distinct genetic architectures that could only be detected with the respective methods, or if they reflect a signal of the same underlying causal locus.

In the original study of (Jacobsson *et al.* 2005), the total effects of all significant and suggestive QTL on BW56 was 634 g, corresponding to 47.3% of the difference between the parental selected lines. Here, the combined effect of the markers retained in the multi-locus model (Table 2) is 527 g. However, when considering only the QTL in the original study that have been replicated here, their originally estimated contribution was 416 g, indicating that the segregation of the QTL alleles in the founder-lines revealed here biased the originally reported estimates that assumed fixation for alternative alleles in the HWS and LWS lines, downwards. The allele-frequencies for the selected markers in the HWS founders show an interesting pattern (Supplementary table 1). Although the weight-increasing alleles are more common in the HWS at all markers, the alternative alleles are still far from fixation in the lines. This suggests that multiple alleles are segregating both within and across the selected lines at the loci with the largest individual effects in the population.

A key for successful imputation of the high-density marker set throughout the AIL pedigree is that the haplotypes across these markers are correctly estimated in the founders. There are several properties of the Virginia-lines that improve haplotype-estimation from high-density genotypes. First, as the number of generations since the lines diverged is relatively few (40 generations), most new haplotypes will result from recombination of original haplotypes, rather than by new mutations. Second, the strong artificial selection imposed on the populations since they were founded is likely to have further reduced haplotype-diversity across the genome. This is likely the reason that many selective-sweeps across long haplotypes have been found to be fixed, or nearly fixed, across the genome within and between the lineages (Johansson *et al.* 2010; Pettersson *et al.* 2013). This is reflected in a large average LD-block size (> 50 kb) across the genome (Marklund and Carlborg 2010). Given the density of the 60k SNP-chip genotyping used here, several markers will be present on each such LD-block and hence improve efficiency in haplotype estimation. Additional genotyping will, however, be necessary in subsequent generations to experimentally confirm the associations to imputed markers reported here.

Genotype data is available for all individuals in the AIL pedigree. The dense marker backbone (~1 marker/cM) from the first genotyping of the AIL (Besnier *et al.* 2011), allow the relatively long haplotypes that are inherited as intact segments from parents to offspring to be efficiently phased, imputed and traced throughout the pedigree for later association analyses.

Here, the association-analysis was performed using a linear model including fixed effects of genotype, sex and AIL generation. Sex and generation were included as both these environmental factors had significant effects on BW56 (Besnier *et al.* 2011). Implementing the model-selection by backward-elimination in a bootstrap-based framework is a way to account for possible effects of population-structure in the AIL that might increase the risk for reporting false positives. However, since the association signals in most cases overlap well with the final marker set resulting from the testing of experiment-wide significant associations, we do not find this to be any cause of great concern in this experiment.

### Conclusion

In conclusion, this study shows that the proposed imputation-based association-mapping strategy, and further model selection by backward-elimination in a bootstrap-based framework, is useful for identifying independent association signals within and across the nine evaluated QTL. The association-peaks were narrower than those obtained in the earlier performed linkage analysis, often highlighting regions down to 2-3 Mb in length allowing the identification of multiple association-signals in several QTL. This suggests that the association-based strategy has higher resolution, as well as provides an improved power to disentangle the effects of multiple linked loci inside QTL, compared to linkage-based fine-mapping. Combining traditional linkage-based approaches to analyze outbred Advanced Intercross populations with imputation-based association mapping approaches might thus be an important and cost-effective approach to improve the efficiency in post-association bioinformatic analyses and functional explorations aiming to identify candidate mutations. A previous candidate-gene study based on the nine QTL fine-mapped here have already reported some interesting mutations in growth related genes (Ahsan *et al.* 2013) overlapping with the association signals reported here. Further bioinformatic investigations of the regions fine-mapped here could potentially reveal new important genes and mutations affecting body-weight in these chicken lines and provide new candidate genes for studying the genetic architecture of metabolic traits in other species, including humans.

## Authors' contributions

ÖC and PBS initiated the study. ÖC designed the project with MA and MB; PBS developed the Virginia chicken lines; PBS (with others) designed, planned and bred the Virginia Advanced Intercross Line; PBS and CFH designed, planned, bred, bled, phenotyped and extracted DNA from the Virginia Advanced Intercross Line population; ÖC designed the statistical analyses; MA, MB and ÖC contributed analysis scripts; ÖC, MB and MA performed the data analyses, summarized the results and wrote the manuscript. All authors read and approved the final manuscript.

## Acknowledgements

Formas (grant 221-2013-450 to ÖC) and the Swedish Research Council (grant 621-2012-4634 to ÖC) are acknowledged for financial support. We thank Leif Andersson for initiating the AIL experiment with PBS and sharing the data from the F2 intercross. Per Wahlberg and Francois Besnier are acknowledged for their valuable contributions during preparation and quality control work of the genotype and phenotype data from the AIL. Genotyping was performed by the SNP&SEQ Technology Platform in Uppsala, which is part of Science for Life Laboratory at Uppsala University and is supported as a national infrastructure by the Swedish Research Council (VR-RFI).

---

**Table S1.**
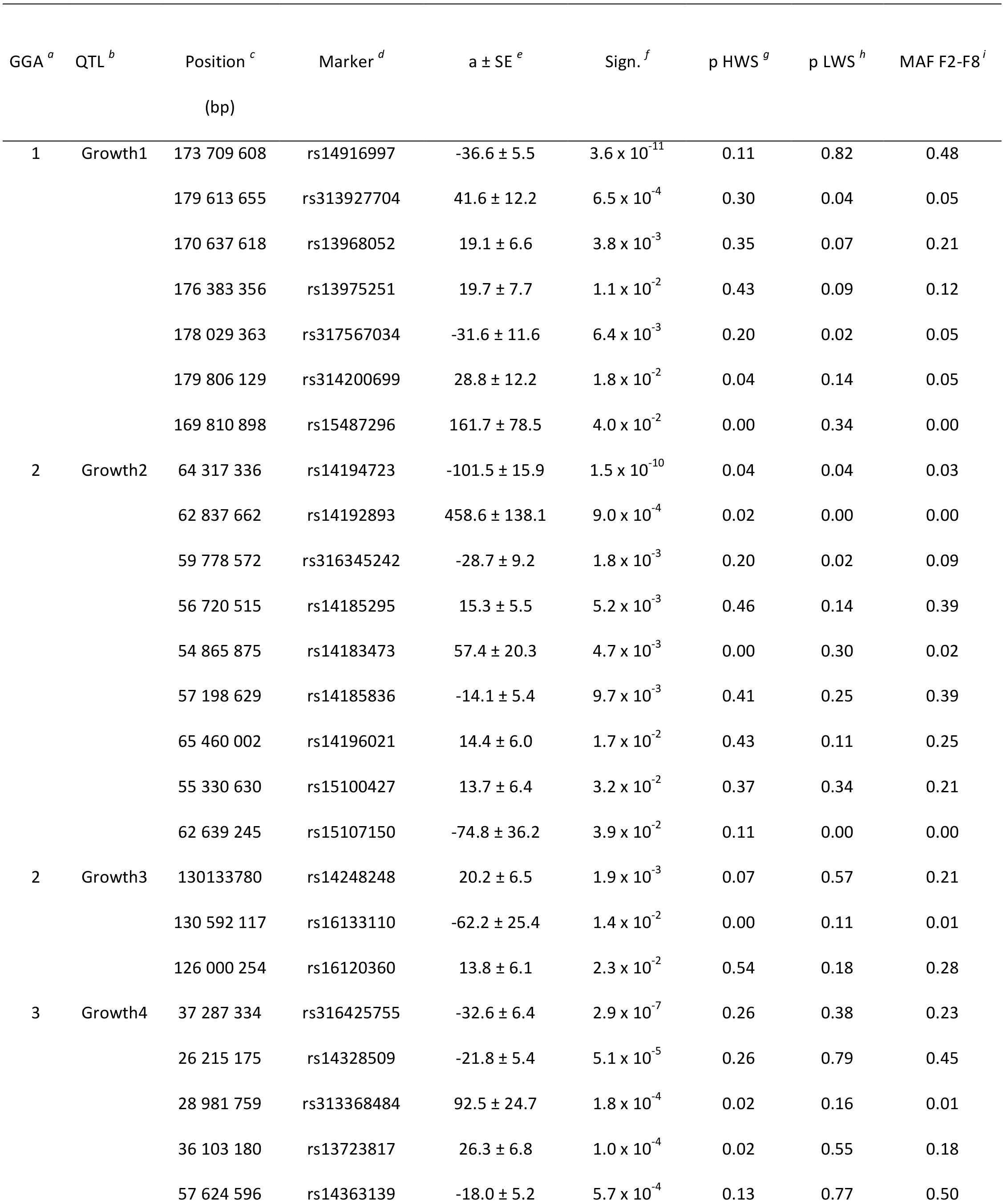

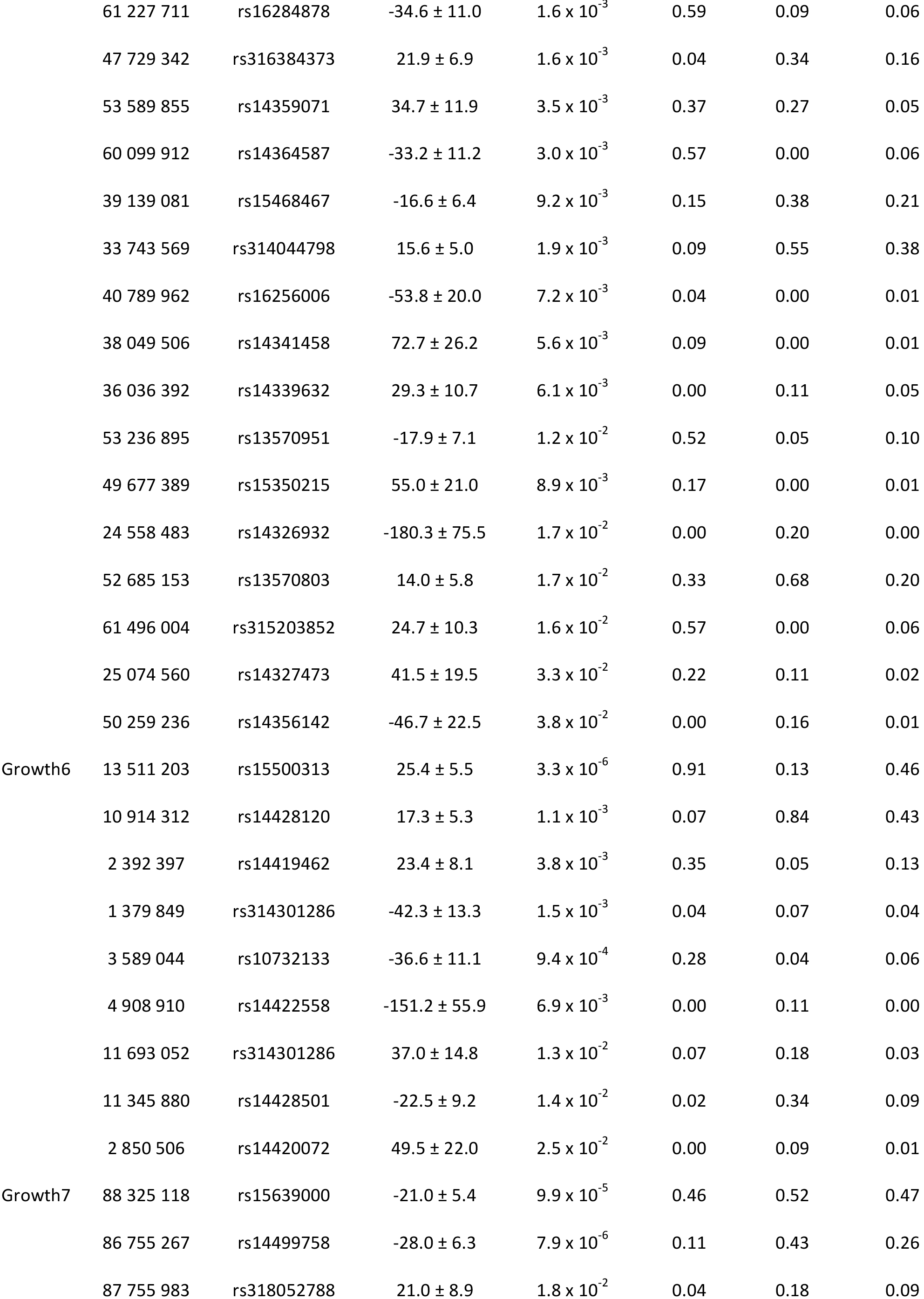

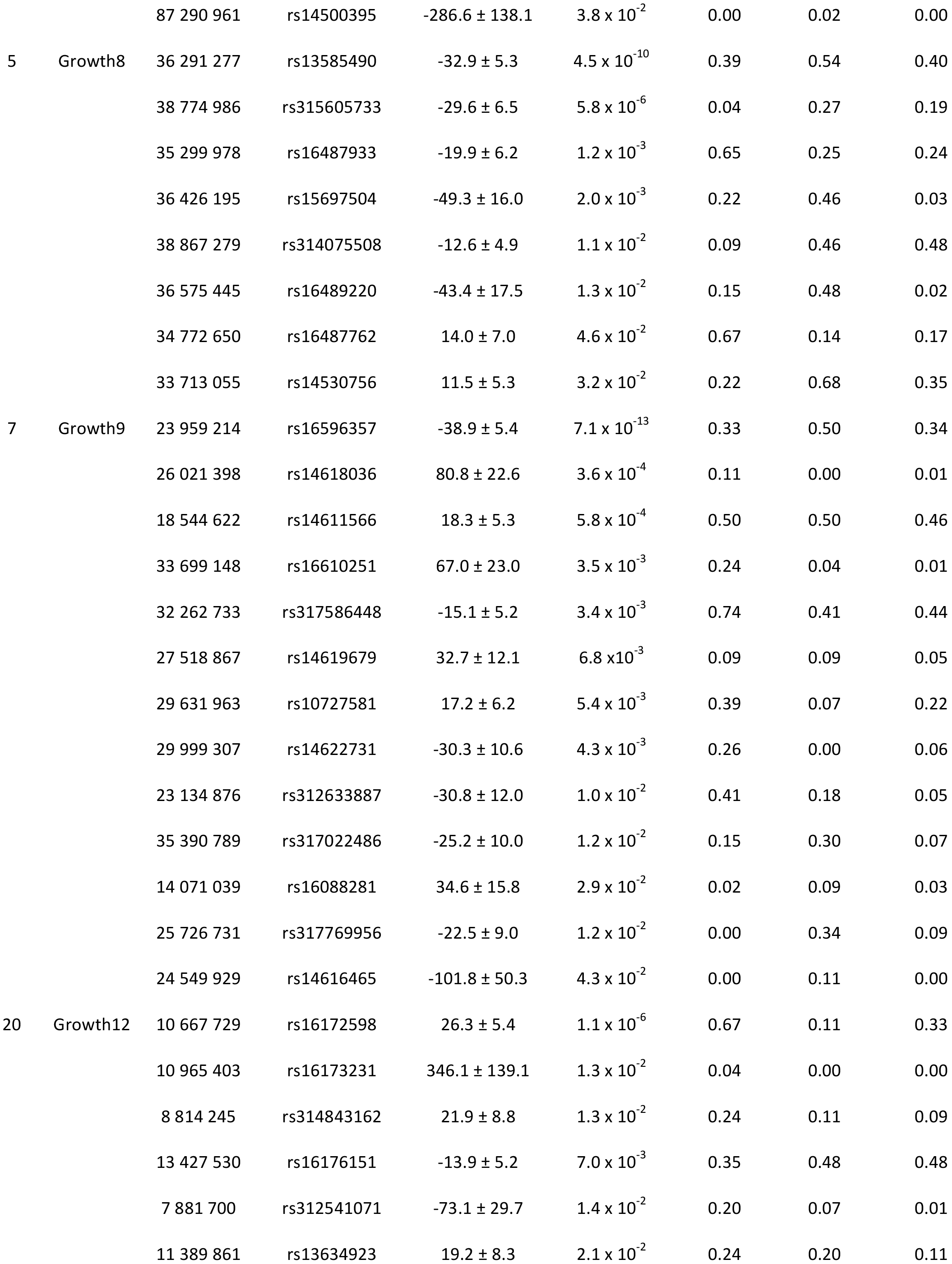

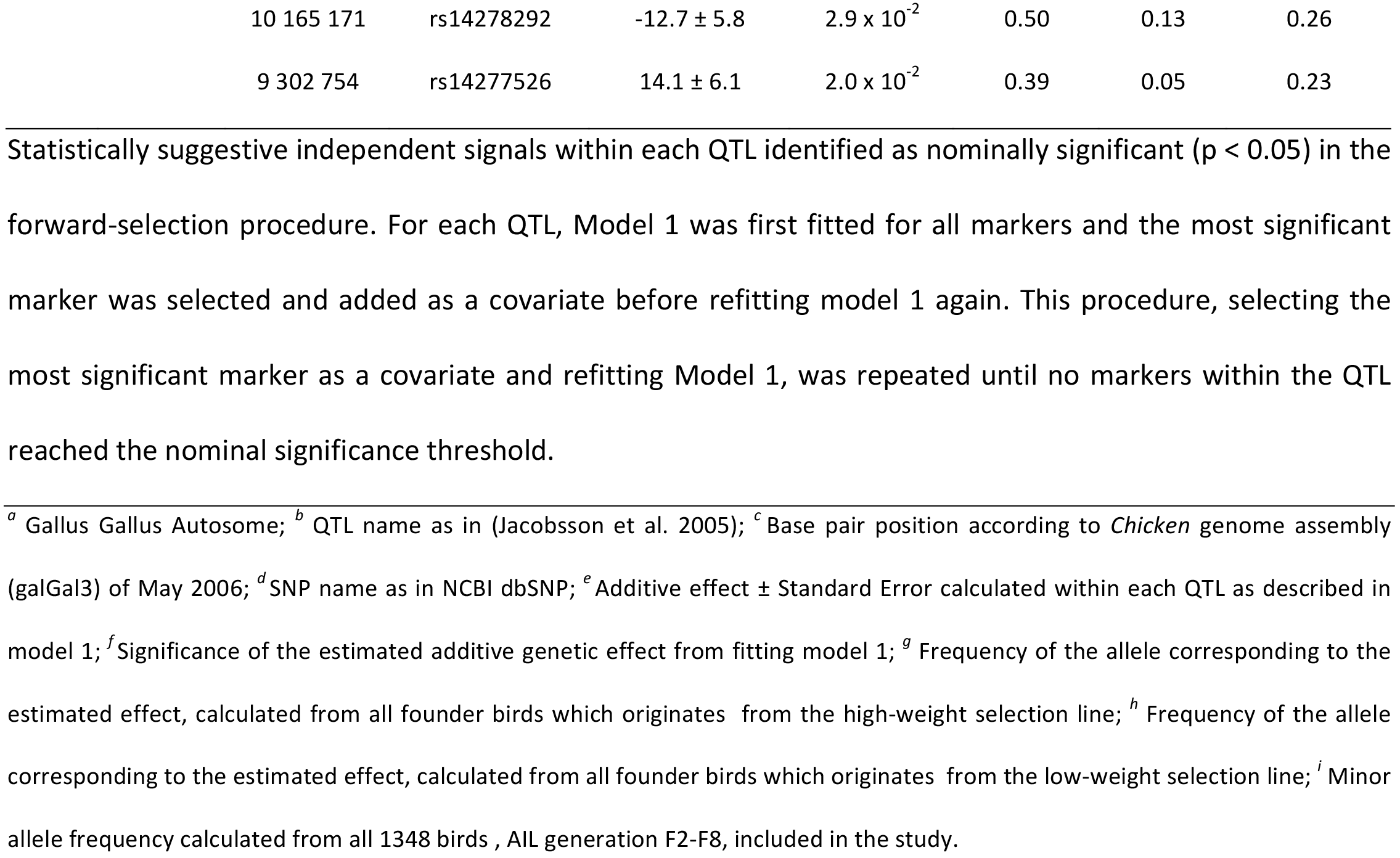

